# Transmission Chains or Independent Solvers? A Comparative Study of Two Collective Problem-Solving Methods

**DOI:** 10.1101/770024

**Authors:** Kyanoush Seyed Yahosseini, Mehdi Moussaïd

## Abstract

Groups can be very successful problem-solvers. This collective achievement crucially depends on how the group is structured, that is, how information flows between members and how individual contributions are merged. Numerous methods have been proposed, which can be divided into two major categories: those that involve an exchange of information between the group members, and those that do not. Here we compare two instances of such methods for solving complex problems: (1) transmission chains, where individuals tackle the problem one after the other, each one building on the solution of the predecessor and (2) groups of independent solvers, where individuals tackle the problem independently, and the best solution found in the group is selected afterwards.

By means of numerical simulations and experimental observations, we show that the best performing method is determined by the interplay between two key factors: the skills of the individuals and the difficulty of the problem. We find that transmission chains are superior either when the problem is rather easy, or when the group is composed of rather unskilled individuals. On the contrary, groups of independent solvers are preferable for harder problems or for groups of rather skillful individuals. Finally, we deepen the comparison by studying the impact of the group size and diversity. Our research stresses that efficient collective problem-solving requires a good matching between the nature of the problem and the structure of the group.

## 1 Introduction

Collective problem-solving and the related concepts of swarm intelligence and collective intelligence have been studied in a wide variety of domains. In biological systems, examples include the nest construction in eusocial insects (Seeley and Visscher, 2004; Khuong et al., 2016) or collective foraging in group-living species (Couzin, 2009). In robotics and artificial intelligence, swarms of relatively simple agents can explore and solve optimization problems efficiently (Garnier et al., 2007; Rendell et al., 2010). Likewise, humans can solve problems in groups during discussions (Kerr and Tindale, 2004), by means of wisdom of crowds procedures (Herzog et al., 2019), or when creating Wikipedia articles (Yasseri et al., 2012; Malone, Laubacher, et al., 2010). Despite this considerable diversity of examples and application domains, all instances of collective problem-solving come down to one central challenge: How should the group be structured to produce the best possible collective output?

Numerous procedures have been proposed to that end. These can be divided into two major categories (Koriat, 2015; Moussaïd, Garnier, et al., 2009): (1) those that involve an exchange of information between the group members, and (2) those that do not. In the first category, direct or indirect interactions among individuals can lead to the emergence of a collective solution (Woolley, Chabris, et al., 2010; Malone and M. S. Bernstein, 2015; Heylighen, 2016). With direct interactions group members exchange information directly via physical signals. Group-living animals, for example, communicate by means of acoustic and visual cues to detect and avoid predators (Domenici et al., 2014; Couzin et al., 2005; Ward et al., 2008). In human groups, the most common case of direct interaction for solving problems takes the form of group discussions (Stasser and Stewart, 1992; Stasser, 1985), where all group members can freely share ideas and strategies to tackle a problem. This approach can produce good results (Toyokawa et al., 2019; Woolley, Aggarwal, et al., 2015; Trouche et al., 2014; Baron, 2005), but is also subject to numerous detrimental effects such as opinion herding, groupthink, and the hidden profile effect (Janis, 1972; Stasser, 1985; Stasser and Stewart, 1992). Also, direct interaction in humans becomes difficult to apply when groups are too large, when members do not work at the same time, or when they have no easy means of communication (e.g., interactions between algorithms and humans (Crandall et al., 2018)).

Some of these limitations can be overcome by *indirect* interaction (Moussaïd and Yahosseini, 2016). In this case, individuals are not directly in contact with one another, but work separately on a common shared group solution. This type of interaction (also known as stigmergy in biological systems) has been heavily investigated in social insects (Heylighen, 2016; Theraulaz and Bonabeau, 1999). For example, when ants engage in the construction of a nest, individuals adapt their behaviour to the current state of the collective construction, which reflects the cumulative actions of all other ants (Khuong et al., 2016). Information is therefore exchanged indirectly, via the collective solution, with no need for direct communication between individuals. This principle can also be applied human groups; a Wikipedia article, for example, emerges mostly as the result of indirect interactions between multiple contributors (Malone, Laubacher, et al., 2010; Heylighen, 2016). In the simplest case, indirect interaction takes the form of a transmission chain, where group members work on a problem sequentially, one after another (Mesoudi and Whiten, 2008). Each individual starts from the final solution of her predecessor and try to improve it, hence gradually giving rise to a collective solution that accumulates the contributions of all group members. Transmission chains have traditionally been investigated in the context of cultural evolution (Mesoudi and Whiten, 2008; Henrich and Boyd, 1998; Kirby et al., 2008; Tomasello et al., 1993) and have more recently been applied to other domains (Moussaïd, Brighton, et al., 2015; Moussaïd and Yahosseini, 2016; Derex and Boyd, 2015).

Beside all these methods, there exists a second class of approaches that do not involve any form of information transfer between individuals. In these cases, individuals first solve the same problem independently and in isolation, and their solutions are eventually combined by an external entity to produce the collective outcome (Koriat, 2015). The most prominent example of such procedures is the *wisdom of crowds*, in which individual solutions are merged by means of statistical aggregation function, such as the mean or the median of all solutions (Herzog et al., 2019). Wisdom of crowds methods are easily scalable as they allow for arbitrarily large group sizes and can yield to accurate solutions (Mannes et al., 2014; Herzog et al., 2019; Surowiecki, 2004). One drawback, however, is that most statistical aggregation techniques cannot be easily applied to complex, multi-dimensional solutions, such as when optimizing a protein folding configuration (Cooper et al., 2010; Romero et al., 2013), improving quantum transport techniques (Sørensen et al., 2016), or trying to solve a jigsaw puzzle (Kempe and Mesoudi, 2014). Thus for problems that have a complex solution structure, the most common practice consists in collecting a large number of independent and hence diverse solutions and choosing the best one at the end (Cooper et al., 2010; Sørensen et al., 2016).

In this work, we specifically focus on problems that have such a multidimensional solution structures. How should a group be structured in this case? In particular, we compare two types of methods: groups that work on the collective solution sequentially, such as transmission chains, or groups that work on independent solutions in parallel? Consider, for instance, the traveling salesman problem – an optimisation task where one has to find the shortest path connecting all cities on a map exactly once (Ochoa and Veerapen, 2018). In a transmission chain, the first individual proposes her solution and transfers it to the next person, who will try to optimize it and pass it in turn to the third one, and so forth. This process continues until all individuals of the group have worked on the collective solution. Would the emerging collective solution be better or worse than when letting all group members search independently and choosing the best one at the end?

In the present paper, we compare the performances of these two methods by means of a behavioural model and a dedicated experiment. In particular, we study how the performances of both methods are influenced by (1) the difficulty of the problem, (2) the skills of the individuals, (3) the group size, and (4) the group’s diversity.

To address these questions, we model problem-solving as a search task (Newell and Simon, 1972; Lazer and Friedman, 2007). We assume that individuals are searching for the best possible solution in a multi-dimensional NK-landscape representing the solution space (see figure 1 and Kauffman and Levin, 1987). For the transmission chains, individuals search sequentially, one after another, each one starting from the last position of her predecessor. The collective solution is then given by the last person’s final position in the landscape. For independent solvers, all individuals start at the same initial position and search in parallel without interactions. The collective solution is then given by the best final solution of all individuals.

Furthermore, we manipulate two variables: the individual skill *S* and the ruggedness of the landscape *K*. We define *S* as the number of dimensions a given individual is able to manipulate (with *S* ≤ *N*, *N* being the total number of dimensions of the NK-landscape), assuming that more skilled individuals are capable of manipulating more dimensions during their search. For instance, an individual with *S* = 2 searching in a NK-landscape with *N* = 10 can only manipulate two out of the ten dimensions in the landscape. The value of *S* can thus be interpreted as the individual’s ability to solve that specific problem.

**Figure 1:**
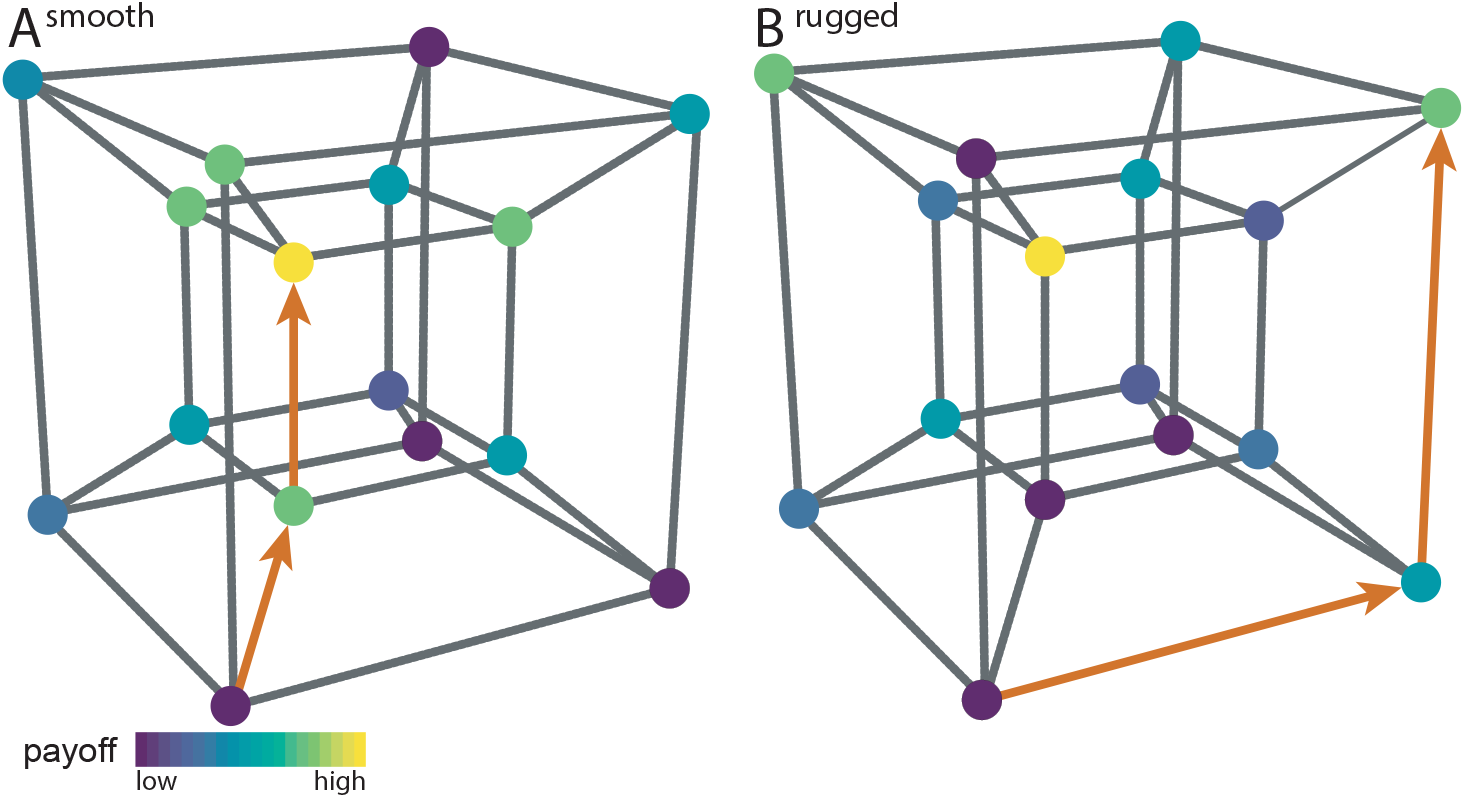
Two examples of four-dimensional NK-landscapes (*N* = 4). Each dot represents one solution and the links between the dots indicate a change along one dimension, illustrating that one can directly move from one solution to another. The color-coding indicates the payoff associated with each solution. In **(A)**, the landscape is smooth (*K* = 1) and has one single locally and globally optimal solution (in yellow). The search trajectory (in red) illustrates how a searcher could reach the optimal solution by gradually moving to the highest neighboring solution. In **(B)**, the landscape is rugged (*K* = 3). It has numerous local optima (i.e. solutions where all neighbouring solutions give a worse payoff) and a simple exploration strategy based on gradual improvements is likely gets stuck on a sub-optimal solution.

The parameter *K* represents the ruggedness of the landscape, i.e. the number of local optima, and is used as a measure of the problem difficulty (see the methods section for more details). Smooth landscapes (with low values of *K*) are reminiscent of problems that are well understood and as such can be easily solved with gradual optimization. In contrast, rugged landscapes (with high values of *K*) have a noisier structure, and can be interpreted as unstructured problems where gradual optimization is usually not an efficient strategy.

## 2 Results

### 2.1 Numerical simulation

We first propose a heuristic model to describe how individuals search in multidimensional landscapes. For this, we assume that individuals randomly manipulate one dimension of their current solution and switch to that new solution if it produces a better payoff than the current one (Gigerenzer et al., 1999; Barkoczi and Galesic, 2016). We use this model to simulate transmission chains and independent solvers, while systematically varying the difficulty of the problem *K*, and the individuals’ skill *S*. As shown in Figure 1A, both methods are influenced by *K* and *S*. As intuitively expected, performances decrease with increasing problem difficulty, and increase with increasing individual skill. However, the transmission chains are less sensitive to the individual skill than the independent solvers, giving rise to two zones of interest as shown in figure 2B: (1) In the lower left corner – for rather easy problems and unskilled individuals – transmission chains outper-form independent solvers, (2) in the upper right corner – for rather difficult problems and skilled individuals – groups of independent solvers perform better. Between these two zones, the performances of both methods become increasingly similar.

**Figure 2:**
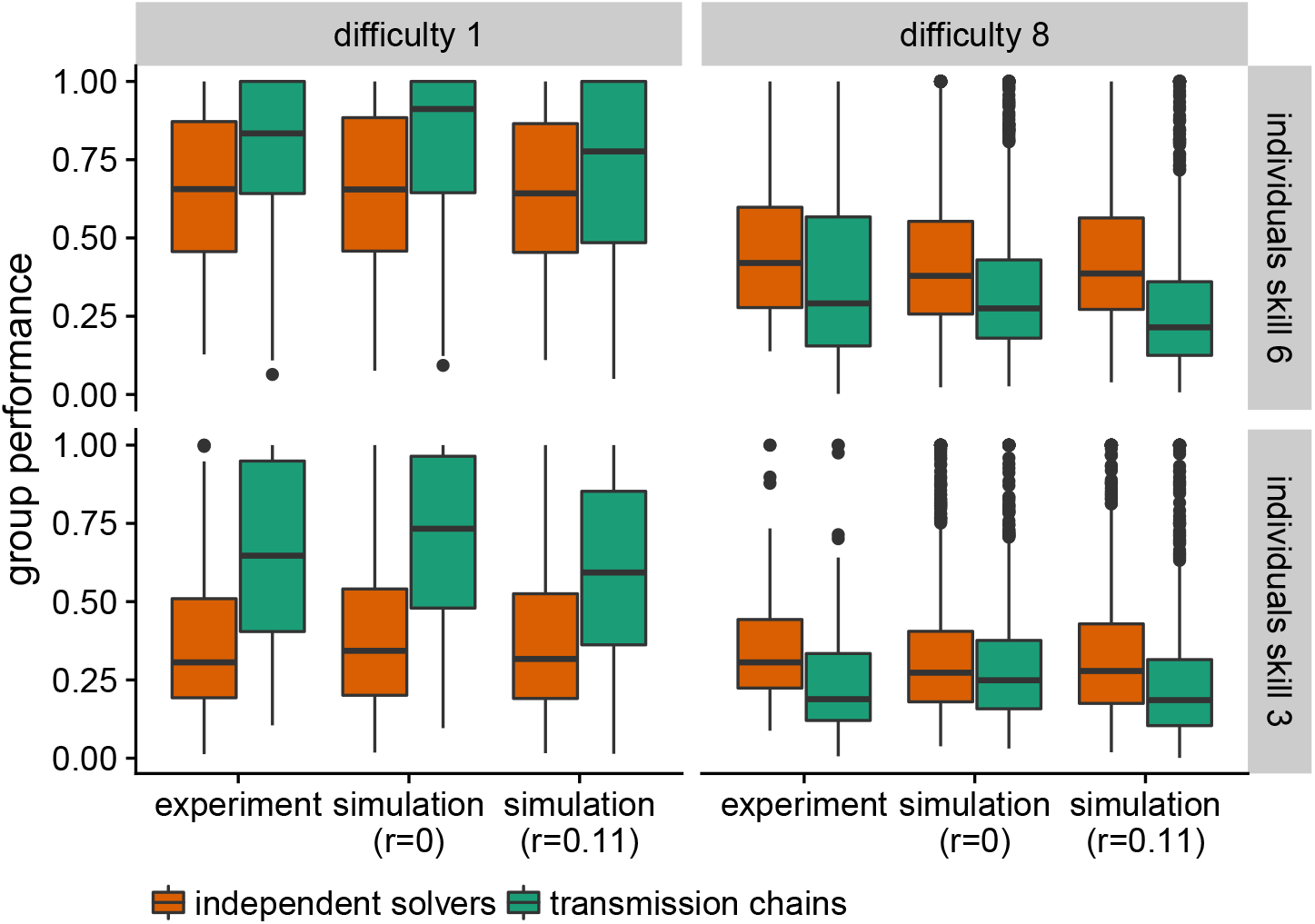
Performance for transmission chains and independent solvers as obtained by numerical simulation, for a group size of 8 individuals and *N* = 10 dimensions. **(A)** Average collective performance of the two methods for varying degrees of difficulty *K* and individual skill *S*. **(B)** Difference in performance between the two methods. Positive values, in blue, indicate that independent solvers outperform the transmission chains and vice versa, in red.

### 2.2 Experimental data

Our simulations suggest that the difficulty of the problem and the individual skills determine which of the two methods performs better. We verify this prediction by means of a controlled experiment. In the experiment, participants searched for the best possible solution either in smooth or rugged landscapes (*K* = 1 or *K* = 8, respectively), and with low or high individual’s skill (*S* = 3 or *S* = 6, respectively). Participants were either part of a transmission chains or in a group of independent solvers (see the Methods section for the detailed procedure). As shown in figure 3, our experimental data confirms the model predictions. In smooth environments, transmission chains outperform independent solvers (*t*(338) = *−*6.56, *p ≤* 0.005, 95ci: *−*258.17 *−* 139.09, BF *>* 100), and this difference is larger for low skills (*t*(166) = *−*7.07, *p* ≤ 0.005, 95ci: *−*361.92 *−* 203.90, BF *>* 100) than for high skills (*t*(172) = *−*2.89, *p ≤* 0.005, 95ci: *−*191.24 *− −*36.11, BF= 15.03). In rugged environments, the opposite is true: Independent solvers outperform transmission chains (*t*(336) = 4.02, *p ≤* 0.005, 95ci: 51.00 *−* 148.43, BF *>* 100). However, contrary to the predictions, the difference of performance is not larger for high skills (*t*(168) = 2.86, *p ≤* 0.005, 95ci: 32.44*−*176.53, BF = 13.98) than for low skills (*t*(168) = 2.93, *p ≤* 0.005, 95ci: 30.56 *−* 156.87, BF = 16.48). In other words, our simulations predicted that the transmission chains would perform better for difficult problems than they actually do. Why is that so?

**Figure 3:**
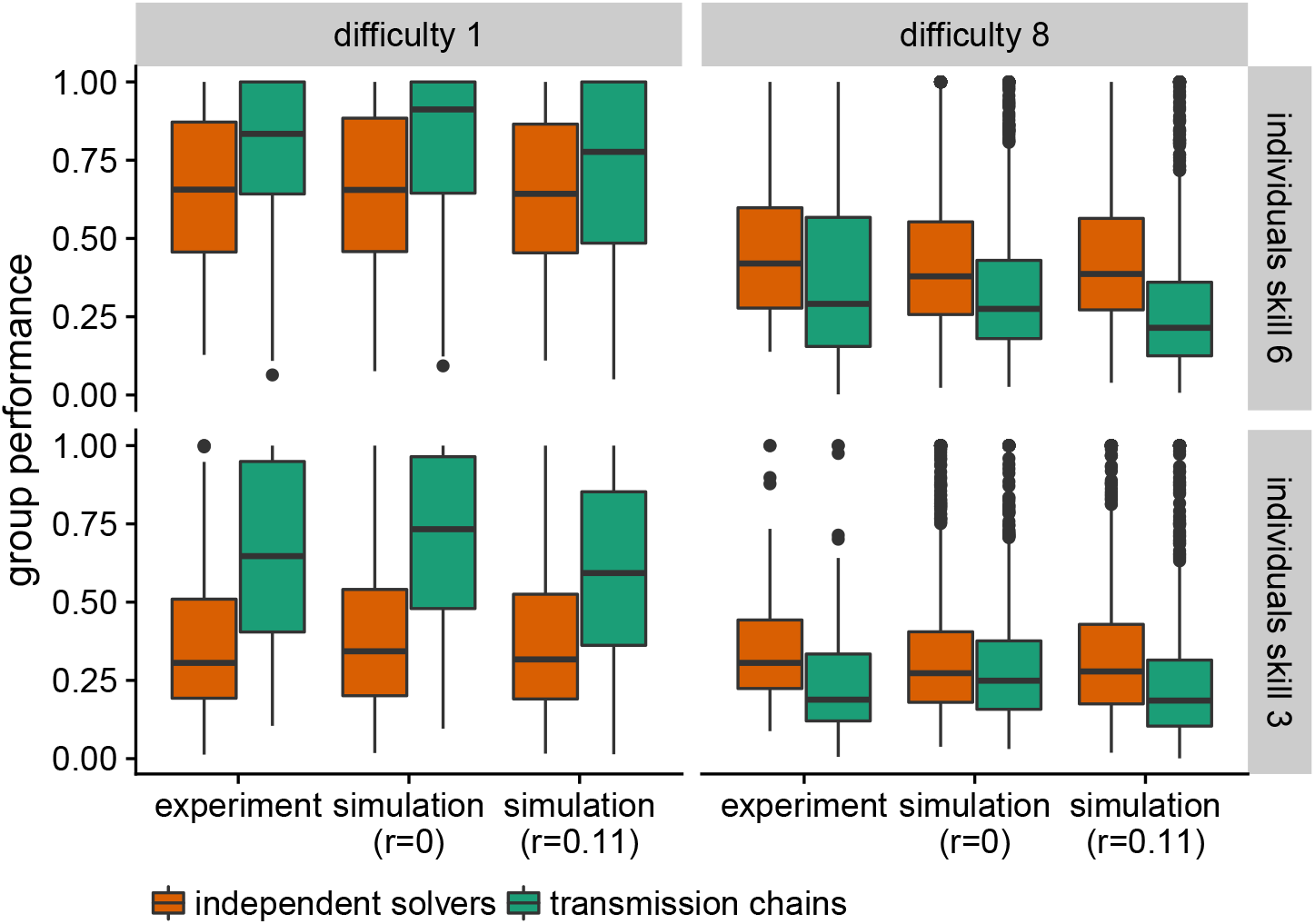
Observed and simulated performances for the transmission chains (in green) and the independent solvers (in orange), for smooth and rugged landscapes (as columns, *K* = 1 and *K* = 8) and for low and high individual skill (as rows, *S* = 3 and *S* = 6). The box plots indicate the interquartile range (box), the median (horizontal line) and 1.5-times interquartile range (whiskers). Outliers are shown as a single dot. For the simulations, the parameter *r* indicates the probability of risky decisions (i.e. the probability to leave a solution for a worse one). The group size is limited to eight individuals.

In transmission chains, performances are considerably affected by the decisions of the last individuals (Moussaïd and Yahosseini, 2016). For instance, the last person of the chain could make the risky decision to leave the good solution found by her predecessors and search for a better one. A failure to do so would impair the collective performance of the entire group.

This behavior is not captured by our search model, in which individuals only leave a solution for a better one (see figure 4). To account for such risky decisions, we extend our model with a parameter *r* reflecting the probability that an individual would not immediately return to the previous solution after sampling a worse one. A positive value of *r* confers some flexibility to the behavior of the agents, preventing them to get stuck at only locally optimal solution, but at the same time increases the risk to lose track of the previous solution. We fit *r* to our experimental data by measuring the frequency of risky decisions in the experimental data (*r* = 0.11). The new simulations match the observed performance very closely (see figure 3, and 4). That is, *r >* 0 decreases the performance of transmission chains for high difficulty and low skills, while slightly improving the performance of groups of independent solvers.

**Figure 4:**
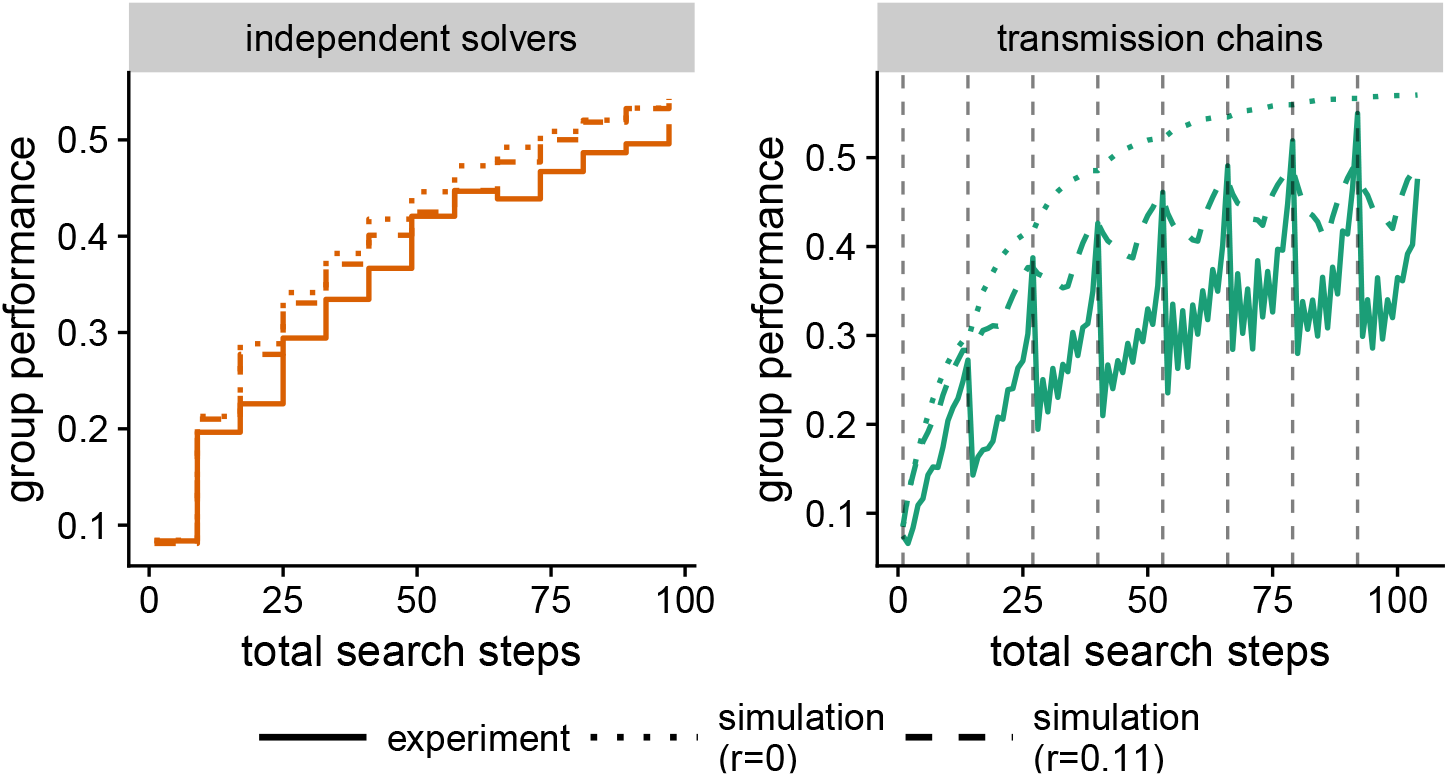
Group performance as a function of total search steps for *S* = 6 and *K* = 1 or 8. Total search steps are the sum of all search steps at the disposal of the entire group. For the transmission chain, a dashed vertical lines indicates that a new individual has started.

### 2.3 Group size and diversity

Finally, we investigate the influence of the group size for the two methods. To account for smaller groups in the experimental data, we simply excluded later individuals to match the desired size and recalculated the group’s performances. Overall, simulations and experimental data exhibit very similar tendencies (see figure 5). In either case, our previous findings are robust to group size variations: with at least three group members, transmission chains outperform independent solvers in smooth environments and are out-performed in rugged ones. In general, group performance increases with group size while the difference between the two procedures remains about the same. The only exception concerns smooth landscapes with high skilled individuals, where both strategies lead a nearly optimal performance.

**Figure 5:**
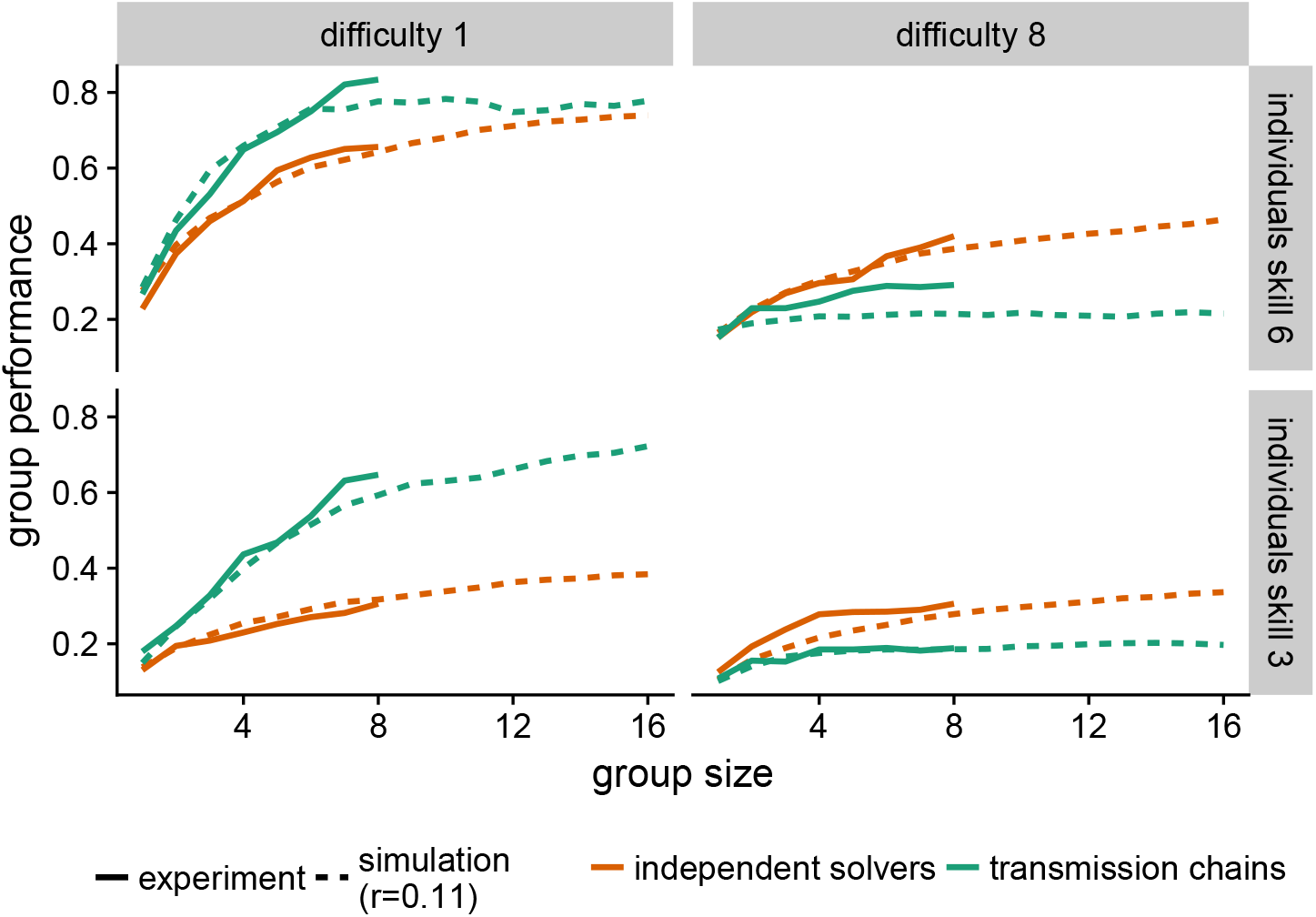
Influence of group size on performance for the two procedures (color-coded), as observed in the experiment data and obtained in simulations.

Overall, we find a diminishing returns for larger groups. In other words, performance does not improve linearly with group size, but eventually plateaus. For example, over all conditions an increase in group size from two to five members improves performance by an average of 0.1 in the experimental data, whereas the same change from five to eight members only improves performance by 0.07. In transmission chains, too many members can even decrease the collective performance, because longer chains increase the risk of losing a good solution due to a risky decision (as described by the parameter *r*).

Finally, we study how the collective performance is impacted by the group diversity — a factor that is known to be critical for collective intelligence (Derex, Perreault, et al., 2018; Jönsson et al., 2015; Tump et al., 2018; Surowiecki, 2004). For this, we define a group’s diversity as the dissimilarity between the dimensions that the group members can manipulate. Put differently, a diverse group is made of individuals that have different perspectives on the same problem. Formally, diversity is measured as the average number of dimensions that only one individual can manipulate in all possible pairs of individuals in the group.

As shown in figure 6, our results reveal very strong evidence for a positive influence of diversity on the performance of independent solvers (*F* (1, 346) = 39.15, BF *>* 100, *adj. r*^2^ = .10) and a moderate one for transmission chains (*F* (1, 343) = 5.47, *BF* = 3.14, *adj. r*^2^ = .01). In short, diversity is indeed a positive factor, and more diverse groups outperform less diverse ones, irrespective of the chosen method.

**Figure 6:**
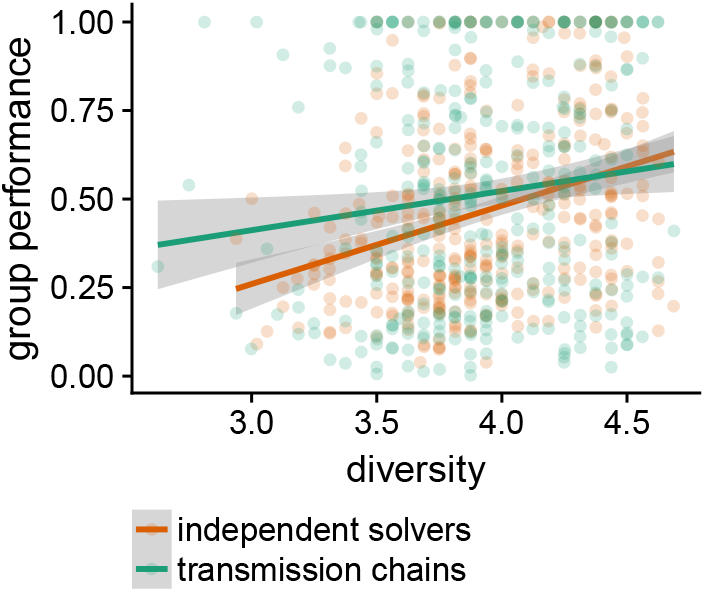
Influence of diversity on the collective performance, as observed in our experimental data. Diversity is defined as the dissimilarity between the dimensions that the group members can manipulate. Each point corresponds to one landscape in one condition. The straight lines indicate the best fitted linear models and the standard error to the two methods (color-coded).

## 3 Discussion

How should a group be structured for solving multidimensional problems? Here we compared two approaches: (1) transmission chains, where individuals tackle the problem sequentially, one after the other, each one building on the solution of its predecessor, and (2) groups of independent solvers, where individuals tackle the problem in parallel without influence and the best solution found in the group is selected afterwards. Our results suggest that the performances of the two methods depend on the interplay between two factors: the problem difficulty (i.e., the ruggedness of the landscape) and the skills of the individuals. Transmission chains outperform groups of independent solvers for easier problems or when the individuals are rather unskilled. In opposition, independent solvers have better performances for difficult problems or when the group members are rather skilled. To put it differently, when trying to continuously improve a solution to a well understood problem or when dealing with rather unskilled individuals, reliance on previous solutions is beneficial. When trying to come up with solutions to an unstructured or ill defined problem in a group of experts one should rather select amongst multiple independent suggestions.

The intuition underlying these results is the following. In smooth landscapes (i.e., easier problems), the global maximum can be found by means of a simple hill-climbing strategy that operates on all the problem’s dimensions. However, low-skilled individuals have only access to a subset of these dimensions. For that reason, independent solvers performs poorly in this case (corresponding to the lower-left corner of the map, figure 2B). Transmission chains, in contrary, combine the skills of the group members. The different dimensions of the problem can therefore be optimized sequentially, explaining the better performance of this method here. As the group members become more skilled (i.e., moving towards the upper-left corner of the map, figure 2B), the difference between the two methods decreases. The collective outcome becomes naturally less sensitive to the chosen method when easy problems are addressed by skillful individuals.

As the problem becomes more difficult, the performances of both methods decline, but the decrease is less pronounced for groups of independent solvers. The challenge of such rugged environments, is that the hill-climbing strategy gets easily stuck at a local optimum. To overcome this issue, individuals need to temporarily reduce their payoff in order to leave the local optimum and explore another region of the landscape. Participants, however, are reluctant to do so (as indicated by the relatively low rate of risky decisions *r* = 0.11). In this situation, independent searchers exhibit better performances because individuals in such groups try different trajectories and arrive at different solutions – thus maximizing the likelihood that at least one of them reaches a good solution. Along these lines, citizen science projects, where people try to solve extremely complex optimization problems, have shown that high-performing participants are less efficient when first exposed to example solutions than when they work independently (Sørensen et al., 2016; Cooper et al., 2010).

Two behavioral factors are driving our results. The first one is the reluctance of individuals to temporarily decrease their payoff, although this is often the only way to find the global optimum. This behavior can be interpreted as risk aversion (Frey et al., 2017), because abandoning a reasonably good solution to search for a better one has considerable chances to fail (Boyd and Richerson, 1996; Miu et al., 2018). Furthermore, research has shown that the willingness to take risks tends to decreases when the problem space becomes vaster and more complex (Yahosseini and Moussaïd, 2019; Mehlhorn et al., 2015), which prevents individuals from “getting lost” in very large problem spaces. In situations where the individuals search is not predominantly guided by payoff (e.g., when individuals are more likely to take risky decisions and move away from a local optimum, or when the pay-off information is not immediately available), additional simulations indicate that transmission chains would outperform independent searchers (as shown in figure 8 in the appendix).

The second important behavioural factor is related to the payoff structure. In our setup, individuals are only rewarded for their personal payoff, but not for the group’s performance. This incites individuals to follow rather conservative search strategies and avoid the risk of losing a good solution. Adding a “safety net” – for instance by rewarding the collective performance – could possibly increase the frequency of riskier search behaviors, favoring the discovery of *leaps* – truly novel and substantially improved solutions (Miu et al., 2018).

In our simple implementation of transmission chains, the collective performance depends substantially on the last individuals of the chain. In other words, the system has no memory of the past solutions, which can result in losing track of a very good solution (Moussaïd and Yahosseini, 2016). This leads to very volatile collective performances over time, as observed in our experimental data and when introducing risky decisions in the simulations (see figure 4). This is along the lines of previous research showing that less inclusive strategies, i.e. strategies that depend on a smaller number of individuals, are more prone to wrong judgments, outliers and noise (Herzog et al., 2019; Mannes et al., 2014). Nevertheless, research in cultural evolution -– studying how an innovation can emerge as solutions are passed from person to person, across generations -– has shown that more sophisticated forms of transmission chains can retain memory of past events and yield more stable collective results.

Future research will consider mixtures and variations of these collective problem-solving methods, such as alternating phases of influenced and independent search (E. Bernstein et al., 2018), or mixing direct and indirect interactions in more elaborated chain structures (Mesoudi and Thornton, 2018).

## 4 Methods

### 4.1 Search environment

The search environments used in our simulations and the experiment were produced by means of the NK-model, which generates multi-dimensional tunably rugged landscapes (Barkoczi and Galesic, 2016; Lazer and Friedman, 2007). The structure of these landscapes is determined by the two eponymous parameters: *N* is the number of binary dimensions and *K* controls the ruggedness by varying the number of interdependencies between each dimension. Low values of *K* generate smooth landscapes with few or no local maxima, which are easy to solve by means of a local optimization procedure (i.e. hill climbing). In contrast, high values of *K* create rugged landscapes with many local maxima, where local optimization is not an efficient search strategy (see figure 1 for a visualization of two NK-landscapes and (Kauffman and Levin, 1987) for a more detailed description of the underlying model).

The NK-landscapes used in our study were generated by fixing *N* = 10 (i.e. our landscapes have 10 binary dimensions, corresponding to 2^10^ = 1024 different solutions) and with various values of *K*. Following several authors, we normalized the payoffs in each landscape by dividing them by the maximal achievable payoff and using a monotonic transformation to raise each payoff to the power of eight (Lazer and Friedman, 2007; Barkoczi and Galesic, 2016). This process causes most solutions to be mediocre and only few to be very good.

### 4.2 Simulation procedure

Following Barkoczi and Galesic (2016), we use a minimalistic heuristic model of individual search (which nevertheless captures experimental data surprisingly well). The model assumes that each agent manipulates one randomly selected dimension at a time, and moves to the new solution if it offers a better payoff than the current one. The agent repeats this search behaviour until the end of the search time. We vary the individual’s skill *S* by allowing only *S* randomly selected dimensions to be manipulated by the agent. That is for *S* = 2 the agent can only manipulate two dimensions of the search environment.

The duration of the search is set to 16 consecutive decisions (but our results seem robust to variations of this number) and all results are averaged over 1000 repetitions.

### 4.3 Experimental treatment

Participants were instructed to search for the best possible solution in a NK-landscape. To facilitate the visual representation, all payoffs were multiplied by 1000, and the 10 dimensions of the landscape were represented as 10 light bulbs that could be either on or off (representing the binary values ‘0’ or ‘1’, see the Supplementary figure 7). Not all light bulbs could be manipulated, due to the restrictions imposed by *S* (those were visually marked by a cross). In each round participant could change the state of one light bulb. After their decision, they were informed about their new payoff and where allowed to return to their previous solution before a new round started.

The eight experimental conditions where selected based on preliminary simulation results, and where matched to the four corners of the figure 2B. The eight conditions consisted of smooth and rugged landscapes (*K* = 1 or *K* = 8, respectively), low or high individuals skill (*S* = 3 or *S* = 6, respectively) and transmission chain or independent group. The order of the experimental conditions was randomized. Each participant played a total of 128 levels, that is, 16 landscapes per experimental condition. To prevent participants from searching all possible solutions, the duration of the search was limited to 2 *× S* rounds for all experimental conditions. Groups of eight individuals were formed searching the same landscape in the same condition.

### 4.4 Experimental procedure and participants

Participants were recruited from the Max Planck Institute for Human Development’s pool and gave informed consent to the experiment. The experimental procedure was approved by the Ethics Committee of the Max Planck Institute for Human Development. Participants were first familiarized with the experiment and informed about their goal, the incentives, and the rules of search in the experiment. Figure 7 in the appendix shows the experimental interface.

We invited 50 participants to the behavioural laboratory of the Max Planck Institute for Human Development. Data of two participants had to be excluded due to technical issues. There were 25 females among the remaining 48 participants (mean age = 27.9, *SD* = 5.13). Participants received a flat fee of 8 € plus a monetary bonus based on their total performance (0.16 € per 1000 points, mean bonus = 6.65 €, *SD* = 1.11 €). The average completion time was 33.64 minutes (*SD* = 10.67 minutes).

### 4.5 Statistical tests

All reported t-tests are one sided. We also report Bayes factors (BF), quantifying the likelihood of the data under *H*_1_ relative to the likelihood of the data under *H*_0_. For parametric tests, the data distribution was assumed to be normal, but this was not formally tested (Wu et al., 2018). Our hypotheses also hold for non-parametric wilcoxon ranked sum tests.

## A Supplementary materials

**Figure 7:**
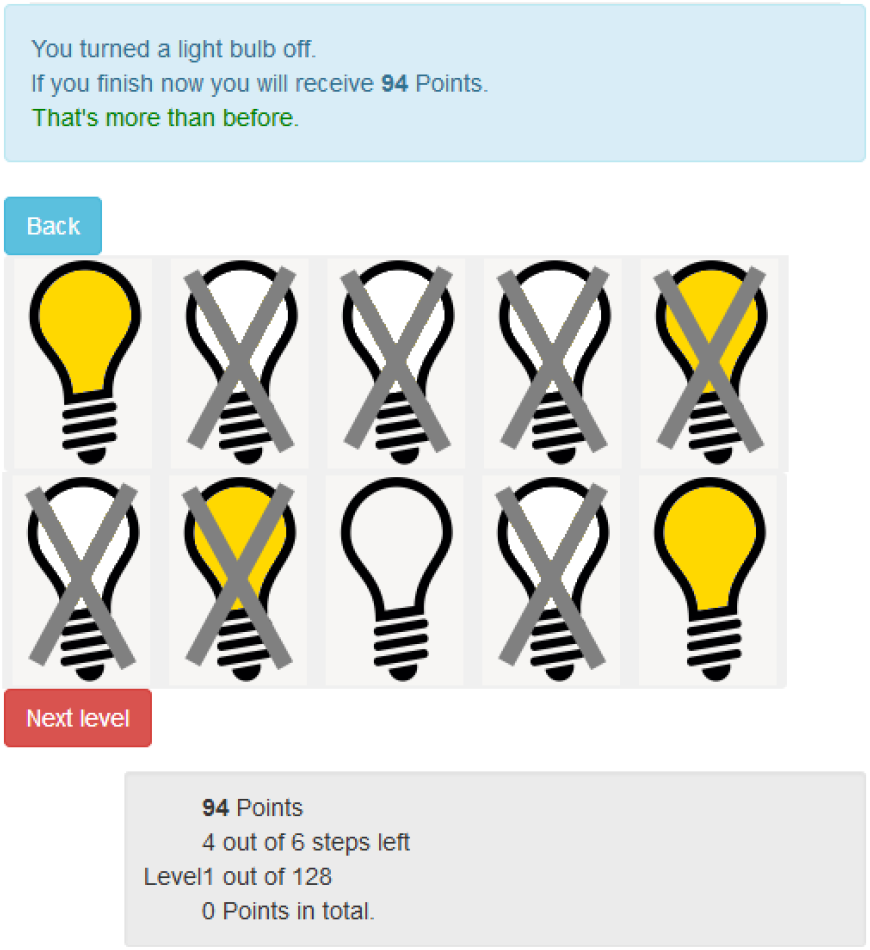
Experimental interface (translated from German to English). The ten light bulbs correspond to the ten dimensions of the the NK-landscape, and the state of each light bubble (‘on’ or ‘off’) represents the values 1 and 0. Grey crosses indicate dimensions that cannot be manipulated due to the restrictions imposed by the individual’s skill *S* (*S* = 3 in this example). Information about the experiment, such as total number of points and remaining number of landscapes (‘level’) are provided in the grey box at the bottom. The blue box at the top shows information related to the previous decision. In each round, participants could either change the state of one light bulb (by clicking on the desired one), skip the remainder of a level if satisfied (by clicking the ‘next level’ button), or return to their previous solution (by clicking the ‘back’ button).

**Figure 8:**
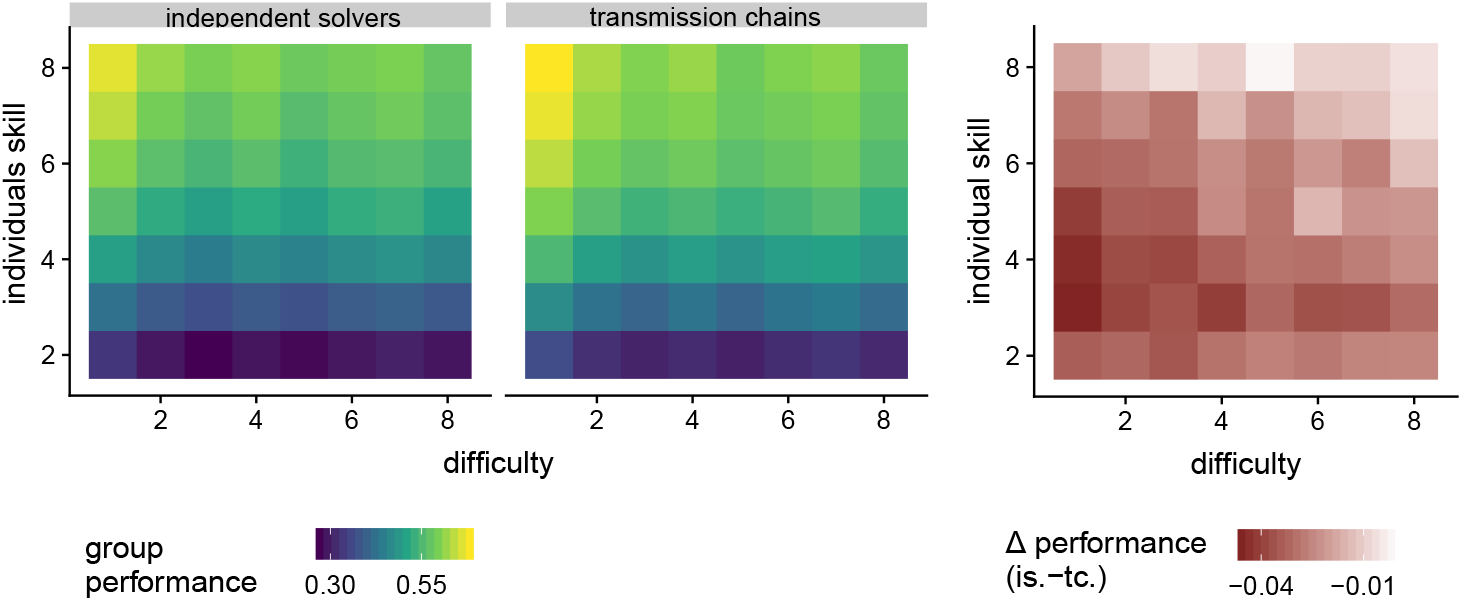
Performance of the transmission chains and groups of independent solvers with a rate of risky decisions *r* = 0.8 and reporting the best solution found. **(A)** Group performance for varying degrees of difficulty *K* and individual skill *S*. **(B)** Difference in performance between the two methods. Positive values indicate that independent groups outperforms the transmission chain, and vice versa.

